# Antibiotic perseverance increases the risk of resistance development

**DOI:** 10.1101/2022.09.20.508757

**Authors:** Gerrit Brandis, Jimmy Larsson, Johan Elf

## Abstract

The rise of antibiotic-resistant bacterial infections poses a global threat. Antibiotic resistance development is generally studied in batch cultures which conceals the heterogeneity in cellular responses. Using single-cell imaging, we studied the growth response of *Escherichia coli* to sub-inhibitory and inhibitory concentrations of nine antibiotics. We found that the heterogeneity in growth increases more than what is expected from growth rate reduction for five out of the nine antibiotics tested. For two antibiotics (rifampicin and nitrofurantoin), we found that sub-populations were able to maintain growth at lethal antibiotic concentrations for up to 10 generations. This perseverance of growth increased the effective population size and led to an up to 40-fold increase in antibiotic resistance development in Gram-negative and Gram-positive species. We conclude that antibiotic perseverance is a common phenomenon across the bacterial kingdom that has the potential to impact antibiotic resistance development.

## Main

The rise in antibiotic-resistant bacterial infections is a global threat to modern medical health care. The dispersal of multi-drug resistant pathogens is driven by the acquisition of chromosomal mutations and mobile genetic elements (*1, 2*). The standard model of resistance development divides selective conditions into ‘sub-MIC’ selection below the minimal inhibitory concentration (MIC) and ‘above-MIC’ selections above the MIC (*3*). Sensitive populations cannot grow during selection above the MIC which makes the existence of pre-existing mutations or resistance elements essential for the development of resistance (*4*). Contrarily, sub-MIC selection conditions permit the growth of sensitive bacteria, and *de-novo* resistance can arise during replication. The novel resistant population often has a growth advantage and can outcompete the sensitive parental population (*5, 6*). Additionally, sub-MIC concentrations of antibiotics can increase conjugation frequencies, select for populations that carry a multi-drug resistance plasmid, and increase bacterial swimming motility as well as biofilm formation (*7-10*). So far, studies on the effect of antibiotics on bacterial growth have generally been performed in batch cultures with large populations thus overlooking any cell-to-cell heterogeneity in the response to antibiotic exposure. Studies using single-cell imaging have shown that resistance factors such as multi-drug efflux pumps are not expressed equally in all cells within a population, which could lead to heterogeneity in the spontaneous mutation rate (*11-13*). Here, we systematically exposed *Escherichia coli* to sub-MIC and MIC concentrations of nine distinct antibiotics and used time-lapse microscopy to analyze the bacterial growth response on a single-cell level. Our main aims were (*i*) to measure cell-to-cell heterogeneity in growth response as a function of antibiotic concentration, and (*ii*) to determine how this heterogeneity affects antibiotic resistance development.

We selected nine distinct antibiotics that cover a wide variety of antibiotic classes and targets, and determined their MIC in Lysogeny broth with 425 mg/L Pluronics (LB; Fig. 1A, Table S1). We then exposed *E. coli* cells to antibiotic concentrations corresponding to 1/8x, 1/4x, 1/2x, and 1x MIC (Table S2) in a modified version of the mother-machine microfluidic chip (*14*) that enabled simultaneous measurements at all four antibiotic concentrations (Fig. 1B). Each microfluidic experiment was divided into three stages: (i) the pre-exposure stage (1 h) where cells were grown in LB to determine the baseline growth rate; (ii) the exposure stage (4 h) where the growth media contained antibiotics; and (iii) the post-exposure stage (3 h) where the growth medium was switched back to antibiotic-free LB to test if the cells were able to recover from the antibiotic exposure (Fig. 1C). Using time-lapse phase-contrast microscopy and automated image analysis, we were able to measure the growth rates of thousands of individual cells during each experiment (Figs. 1D-E, S1-S3). We found that for all nine antibiotics, the cell-to-cell heterogeneity, measured as the relative standard deviation of the growth rates for the growing cells, increased as a function of antibiotic concentration and antibiotic exposure time (Figs. 2, S4). However, only three of the antibiotics (CAR, CRO, and GEN) displayed a significant increase in cell-to-cell growth heterogeneity caused by the specific action of the antibiotics. For the other six antibiotics, the increase in growth heterogeneity could be attributed to slower bacterial growth (Fig. 2, S5). Next, we focused on the last 30 minutes of growth during the 1x MIC exposure phase where the effect of the antibiotics is expected to be the strongest, and analyzed the distribution of growth rates of all cells during this window (Fig. 3). While most antibiotics displayed a unimodal distribution of growth rates around a single average value, nitrofurantoin and rifampicin showed a bimodal distribution. Here, we observed a split between a dead main population and a surviving sub-population that maintained growth with a reduced relative growth rate indicating strong cell-to-cell heterogeneity (Fig. 3F-G).

**Fig. 1.**
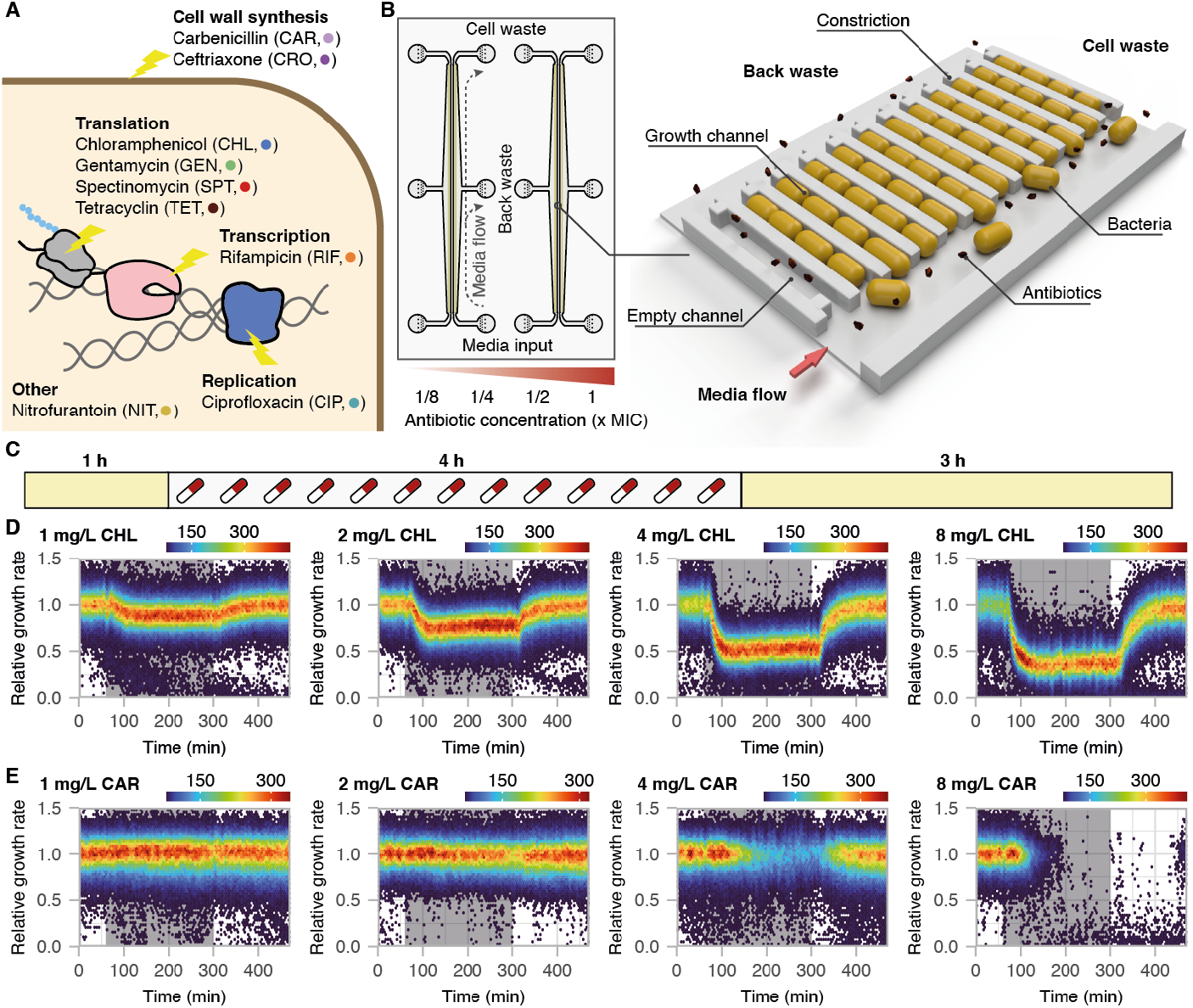
Overview of antibiotic exposure experiment. (A) Antibiotics used in the study. (B) Schematic overview of the microfluidic setup. (C) Timeline of the microfluidic experiments defined by growth in LB (yellow) or LB supplemented with antibiotics (grey). (D, E) Relative growth rates of single cells during the antibiotic exposure experiment. The antibiotic exposure phase is indicated by the grey background. Displayed are representative examples of a bacteriostatic (D) and a bactericidal (E) antibiotic. See figs. S1-S3 for all experiments.

**Fig. 2.**
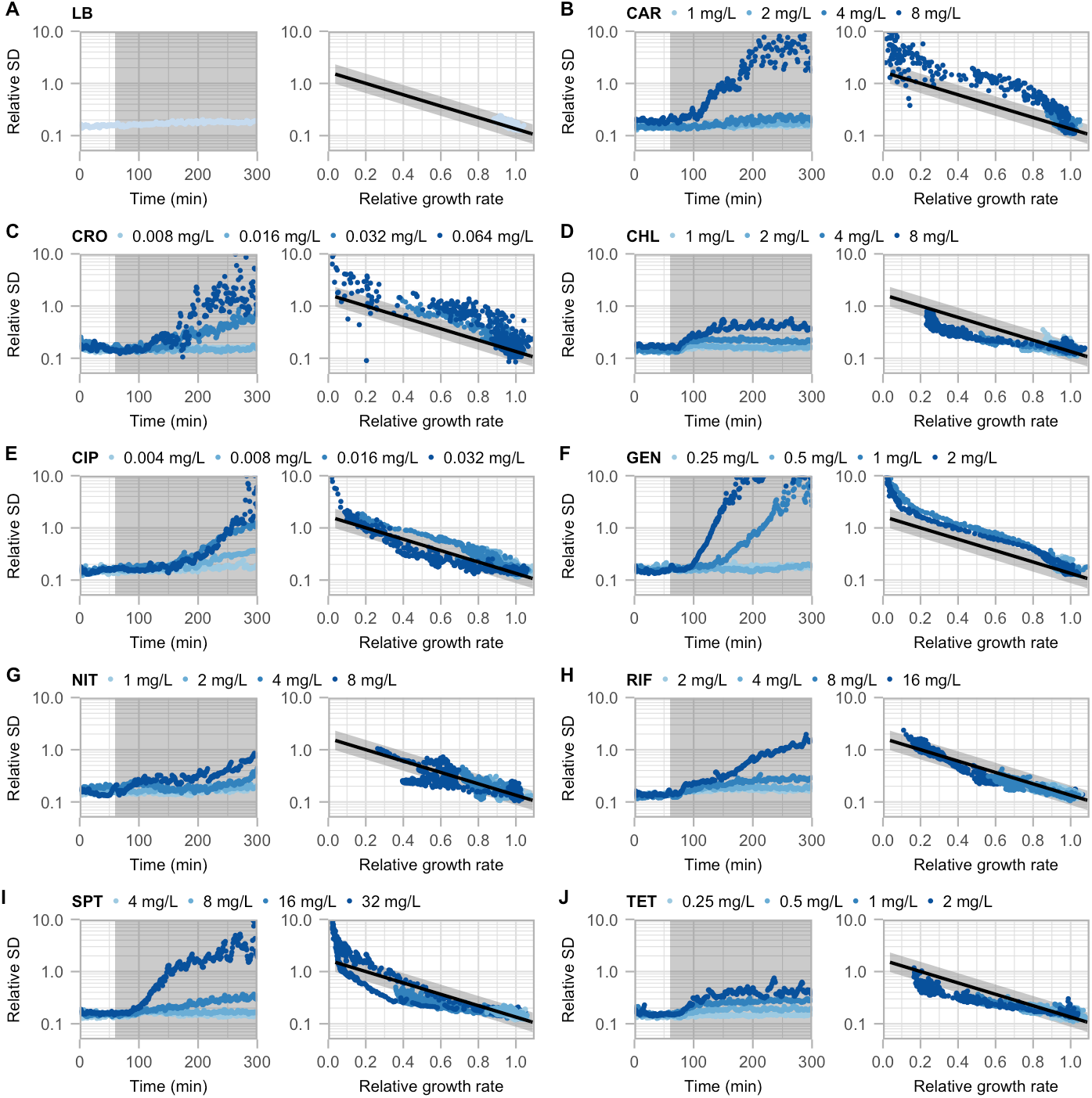
Antibiotic exposure increases cell-to-cell heterogeneity. (A-I) Cell-to-cell heterogeneity, measured as relative standard deviation. (Left) As function of antibiotic exposure time. The period of antibiotic exposure is indicated by the grey background. (Right) As function of relative growth rate. The black line and grey area indicate cell-to-cell heterogeneity that is expected from growth rate reduction (Fig. S8). Depicted are the combined results from two independent experiments. See figs. S4-S5 for the results of each individual experiment.

**Fig. 3.**
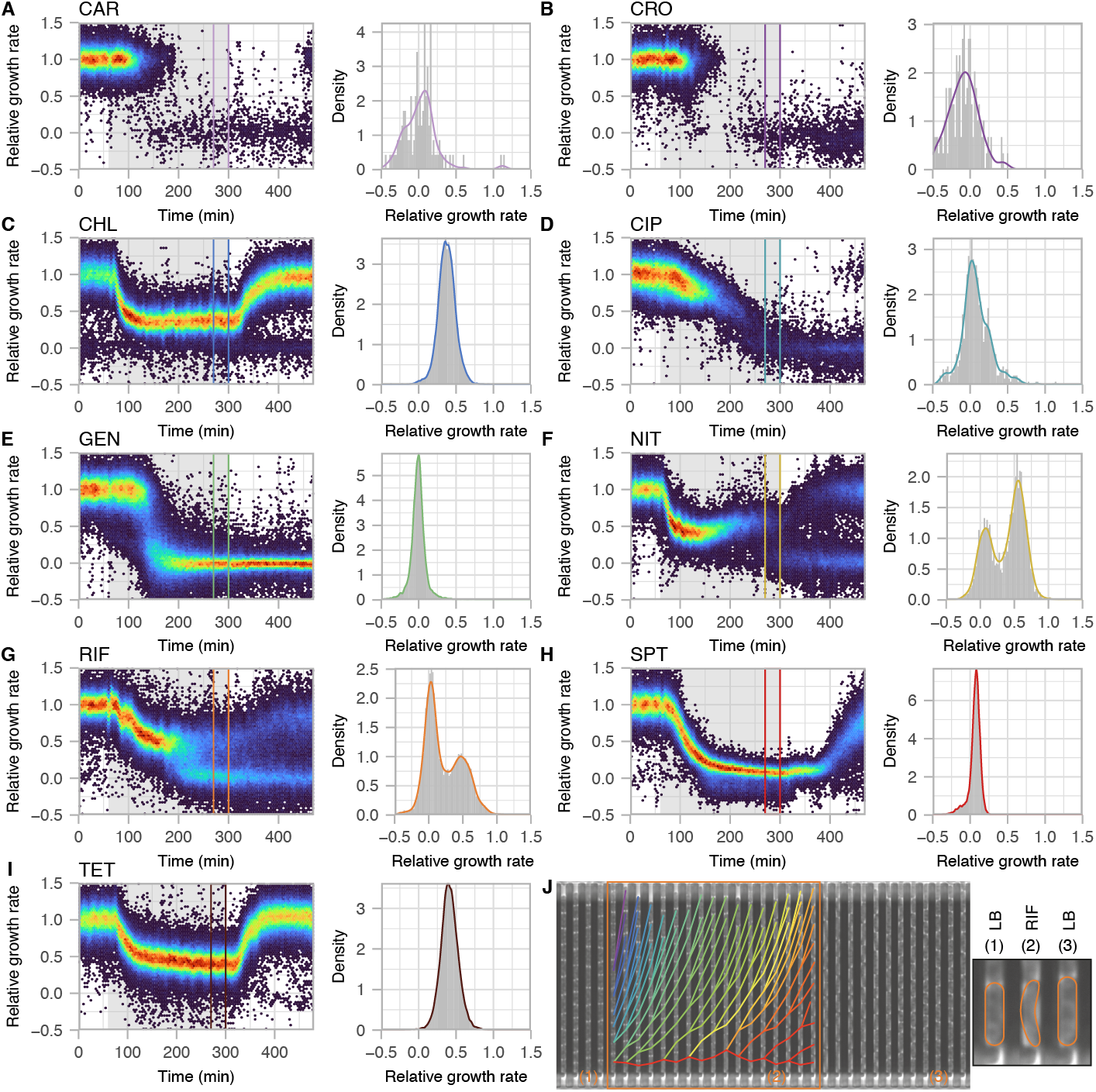
Antibiotic perseverance observed during growth in nitrofurantoin and rifampicin. (A-I, left) Distribution of single-cell growth rates as a function of time during the microfluidic experiments. The antibiotic exposure phase is indicated by the grey background. (right) Density plot of the single-cell growth rates during the last 30 min of antibiotic exposure as indicated by the box in to the left. (J) Time-lapse images (every 15 minutes) of a channel that maintains growth in the experiment displayed in panel G. The phase of antibiotic exposure is indicated by the orange box. The change of position of each cell between images is indicated during the antibiotic exposure phase and branches in the connection lines correspond to cell divisions. The shape of the mother cell during the different phases of the experiment is displayed to the right.

We focused our further analysis on the sub-population that maintained growth in the presence of rifampicin since this antibiotic is commonly used to determine mutation rates (*15, 16*), and the mutations that cause rifampicin resistance are well understood (*17, 18*). A deeper analysis of the growth behavior of the cells during rifampicin exposure showed that the sub-population that maintained growth displayed specific phenotypic changes: (i) the cells were actively dividing with a growth rate of approximately half the growth rate in LB, and (ii) the cell shape changed to slimmer cells. These changes in phenotype were maintained in the daughter cells and fully reverted to the wild-type state within a couple of hours after removal of the antibiotic (Fig. 3G, J). To test if the growing sub-population had an increased MIC, we repeated the experiment but increased the antibiotic exposure time to 17 hours. As before, we could observe the growing sub-population but after approximately 7 h of antibiotic exposure, all growth had ceased, indicating that the sub-population did not have an increased MIC value but rather a reduced rate of growth inhibition (Figs. S6B). We also performed an equivalent experiment with a high concentration of rifampicin (∼6x MIC) and found no growing sub-population, further confirming that no MIC increase is involved in the observed phenotype (Fig. S6C). Due to the lack of MIC increase, we conclude that the observed phenotype cannot be classified as antibiotic resistance or heteroresistance (*19*). Our data show that the cells are actively dividing during antibiotic exposure, which excludes antibiotic persistence, and antibiotic tolerance is a general attribute of the whole bacterial population rather than of a sub-population (*20, 21*). Thus, we conclude that the observed phenotype cannot be properly described by either of these terms and therefore refer to it as antibiotic perseverance within this work.

To quantify the frequency of antibiotic perseverance within the bacterial population, we determined the number of cell divisions of all mother cells within the 17 h exposure experiment (Fig. S6). We found that high-level rifampicin exposure (∼6x MIC) led to a rapid halt of cell division (1.1 ± 0.4 cell divisions). Low-level rifampicin exposure (1x MIC), on the other hand, resulted in a slower termination of growth (3.8 ± 1.2 cell divisions) with a significantly broader cell-to-cell variation based on the standard deviations of the distributions (P < 0.001, two-sided Kolmogorov-Smirnov test) (Fig. 4A). The persevering sub-population represented 2% of the mother cells and displayed a 2.6-fold increase in cell divisions compared to the main population (9.5 ± 0.6 vs. 3.7 ± 0.9 cell divisions). Furthermore, the growth rate of the persevering mother cells was indistinguishable from that of the main population before the antibiotic treatment (P = 0.65, two-sided t-test) (Fig 4B).

**Fig. 4.**
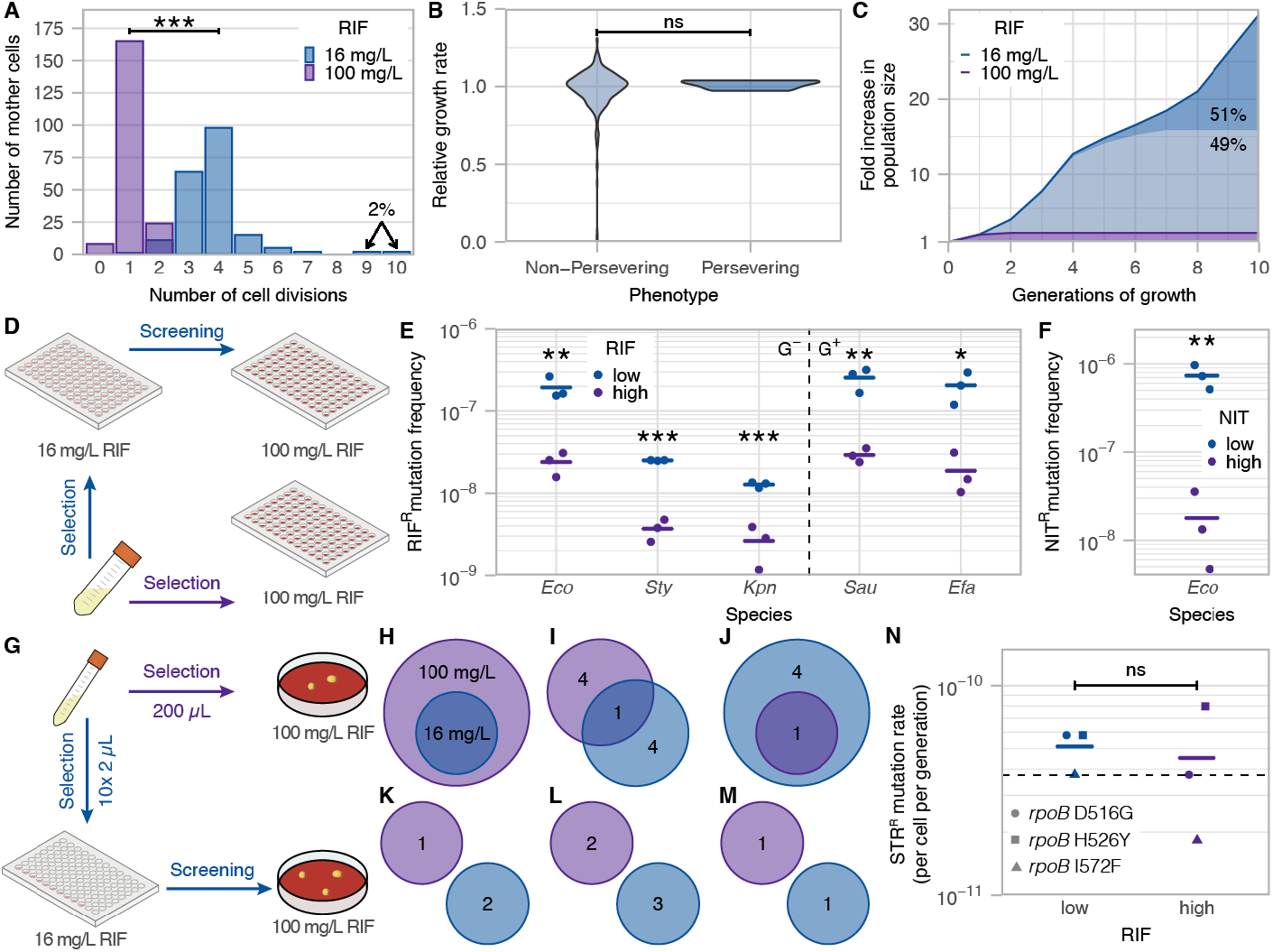
Antibiotic perseverance increases effective population size and frequency of antibiotic resistance development. (A) Distribution of cell divisions of mother cells during the 17 h rifampicin exposure experiment (two-sided KS-test). (B) Violin plot displaying the growth rates of persevering and non-persevering mother cells before antibiotic exposure (two-sided t-test). (C) Modelled change of population size after exposure to rifampicin based on the distribution of cell divisions displayed in panel A. (D) Experimental setup to determine the frequency of high-level rifampicin resistance during different selection conditions. (E, F) Mutation frequencies for high-level rifampicin (E) and nitrofurantoin (F) resistance during selection at low-level and high-level antibiotic concentrations. The line indicates the average value (two-sided t-test). (G) Experimental setup to test if rifampicin resistance mutations are pre-existing in the bacterial culture. (H-M) Venn diagrams for the Null-hypothesis that all rifampicin-resistance mutations are pre-existing within the culture (H) and for rifampicin resistance mutations in the rpoB gene obtained in five independent experiments (H-M). (N) Mutation rate for streptomycin resistance of rifampicin-resistant mutants selected during low and high rifampicin concentrations in the experiments displayed in panels I-M. (ns P > 0.05, * P < 0.05, ** P < 0.01, *** P < 0.001).

To determine the impact of antibiotic perseverance on the bacterial population, we modelled the change in population size after antibiotic exposure at low-level and high-level rifampicin based on the distributions of cell divisions obtained in the 17 h exposure experiment. The total population size increased 14-fold during low-level exposure compared to high-level rifampicin concentrations (31.3-fold vs. 2.2-fold). Importantly, half of the difference (51%) was caused by the 2% of cells that displayed antibiotic perseverance demonstrating that this phenotype can have a significant impact on the bacterial population during antibiotic exposure (Fig. 4C).

Every round of replication can lead to the spontaneous formation of a mutation that could decrease the susceptibility of the resulting strain to the exposed antibiotic (*4, 22*). Thus, sub-populations that maintain growth during lethal selection conditions could increase the risk of antibiotic resistance development. We tested this hypothesis by selecting high-level rifampicin-resistant mutants during exposure to low-level and high-level concentrations of rifampicin and found an 8.3-fold increase in mutation frequency during low-level rifampicin concentrations (P = 0.008, two-sided t-test) (Fig. 4D-E). The increased mutation frequency can be fully accounted for with the 14-fold increase in effective population size during these selection conditions. Next, we tested if the observed mutations indeed appeared after antibiotic exposure, as our model suggested, by analyzing which rifampicin resistance mutations were selected during low-level and high-level antibiotic exposure. The mutation analysis showed that (i) both selection conditions resulted in single amino-acid changes within the *rpoB* gene that encodes the -subunit of RNA polymerase, and (ii) the mutations found during the low-level selection conditions were not pre-existing within the population but must have occurred during the antibiotic exposure time (Fig. 4G-M, table S3). Furthermore, we confirmed that the mutations occurring during low-level rifampicin exposure were not selected in cells with a higher intrinsic mutation rate for example due to inactivation of genes involved in the mismatch repair system (Fig. 4N). These results agree with the proposed model that sustained growth during antibiotic exposure increases the frequency of resistance development.

To test if these results are unique to *E. coli*, we performed a 17 h rifampicin exposure experiment with *Salmonella enterica* serovar Typhimurium and found an equivalent persevering sub-population at low-level rifampicin concentrations (Fig. S7). As before, the mutation frequency for high-level rifampicin resistance was significantly increased (6.8-fold, P = 0.002, two-sided t-test) during exposure to low-level compared to high-level rifampicin concentrations (Fig. 4E, table S4). We also found significant increases in mutation frequency of similar magnitude in all other tested Gram-negative (*Klebsiella pneumoniae*) and Gram-positive (*Enterobacter faecalis, Staphylococcus aureus*) species, indicating that the observed phenomenon is common across the bacterial kingdom (Fig. 4E, table S4). Finally, we tested the mutation frequency for nitrofurantoin resistance in *E. coli* since we also observed a bimodal distribution of growth rates in the presence of nitrofurantoin (Fig. 3F). The results showed that the mutation frequency for nitrofurantoin resistance was increased 40.2-fold when selected at low-level concentration of nitrofurantoin compared to direct selection at the high-level concentration (P = 0.005, two-sided t-test) (Fig. 4F).

## Discussion

Previous research has shown that heteroresistance, persistence, and tolerance can significantly contribute to the development of antibiotic resistance (*19, 23, 24*). Our data describe an additional phenotype that appears to be common among bacteria and applies to at least two antibiotics (rifampicin and nitrofurantoin). The frequency of perseverance (∼2% of cells), the lack of MIC increase, and the reversible phenotype indicate that this phenotype is not caused by genetic changes (*25*). The persevering bacterial cells change growth rate and shape during antibiotic exposure, and this phenotype reverts quickly after the removal of the antibiotics which could indicate a changed state of gene expression within the cells. We therefore propose a model where bacterial sub-populations are prone to become persevering cells due to the intrinsic cell-to-cell variation of gene expression within a bacterial population. This cell-to-cell variability has been demonstrated for multiple resistance-related genes such as efflux pumps and stress response regulators (*12, 26*). The resulting perseverance phenotype enables extended survival and growth during short-term antibiotic exposure (∼7h in our experiments with rifampicin) and increases the risk of antibiotic resistance development. While we have only studied the selection of chromosomal mutations during this growth phase, it is likely that growth under these conditions also increases the risk of conjugational transfer of multi-resistance plasmids or acquisition of resistance genes by transduction or transformation.

## Methods

### Bacterial strains and growth conditions

A list of bacterial strains used in this study can be found in table S5. Bacteria were grown at 37°C in LB broth with surfactant Pluronic F-109 (Sigma-Aldrich 542342, 425 mg/L) or on plates of LB broth solidified with 1.5% agar (LA-plates). Antibiotics were added to the media as described. When indicated, bacteria were grown in defined minimal medium (M9) with surfactant Pluronic F-109 (425 mg/L) and enriched with 0.2% glucose, 0.2% glycerol, and 0.08% RPMI 1640 amino acids (Sigma Aldrich R7131) as described.

### Antibiotics

All antibiotics were ordered from Sigma Aldrich. Stock solutions were prepared in water for carbenicillin (C1389), ceftriaxone (C5793) ciprofloxacin (PHR1044), gentamycin (G1914), and spectinomycin (S4014), in 70% ethanol for chloramphenicol (C0378) and tetracycline (T7660), in methanol for rifampicin (R3501), and in DMSO for Nitrofurantoin (46502).

### Minimal inhibitory concentration

The MICs were determined by broth dilution in 96-well plates. Bacterial colonies were resuspended in 0.9% NaCl to 0.5 McFarland and diluted 100-fold in growth medium. 50 µL cells were added to each well with 50 µL antibiotic solution resulting in a total volume of 100 µL per well containing approximately 105 cfu bacterial cells (106 cfu/mL). The microtiter plates were sealed and incubated for 18 h at 37°C before reading. All MIC tests were performed in three biological replicates. A list of the tested antibiotic concentration ranges and MIC results can be found in table S6.

### Time kill assay

Approximately 106 cfu of exponentially growing cells were added into 2 mL pre-warmed LB with Pluronics containing antibiotics at 8x MIC (∼ 5×105 cfu/mL) and incubated shaking at 37°C. After 0, 1, 2, and 4 h samples were taken, diluted in 0.9% NaCl, and plated on LA plates. Plates were incubated overnight at 37°C after which colonies were counted. Bactericidal activity was defined as >99% reduction of the initial inoculum size.

### Determination of mutation frequency to high-level rifampicin and nitrofurantoin resistance

Overnight cultures were diluted in fresh LB with Pluronics and rifampicin/nitrofurantoin, and aliquots of 100 µL were distributed in the wells of a 96-well microtiter plate. See supplementary tables S4 and S7 for details on the antibiotic concentrations and inoculum sizes for the different species. A dilution series of each overnight culture was plated on LA plated to determine the actual inoculation size for each well. 96-well plates were sealed and incubated for ∼18h shaking at 37°C, and LA plates were incubated for ∼18h at 37°C. From each positive well, 1 µL culture was transferred into 100 µL fresh media containing the high antibiotic concentration to confirm high-level resistance. Plates were sealed and incubated for ∼18h shaking at 37°C before growth was determined. Mutation frequencies were calculated by dividing the number of positive wells by the combined bacterial inoculation size within the 96-well plate. All measurements were performed in biological triplicates and frequencies of resistance development at low and high antibiotic concentrations were compared using a two-sided two-sample t-test.

### Determination of mutation rate to streptomycin resistance

The mutation rate to streptomycin resistance was used to compare the general mutation rates in rifampicin-resistant isolates. Twenty 1 mL cultures of each isolate were grown overnight shaking at 37°C. Each culture was centrifuged for 5 min at 7,000 rpm in a benchtop centrifuge (5424 R, Eppendorf). The bacterial pellets were resuspended in approximately 100 µL media, plated on LA plates containing 100 mg/L streptomycin, and incubated overnight at 37°C. Mutation rates were calculated according to the formula

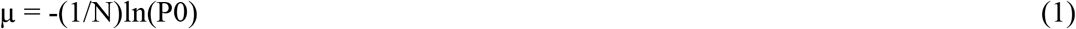

where µ is the mutation rate, N is the number of viable cells per plate (∼2.8 × 109 cfu), and P0 is the proportion of plates that did not give rise to streptomycin-resistant colonies. A Chi-squared test based on the number of cultures with and without streptomycin-resistant colonies was used to compare the results from the rifampicin-resistant isolates with wild-type E. coli. The results of the assay are shown in table S8.

### Test for pre-existing mutations

Five independent cultures of 1 mL LB with Pluronics were inoculated and grown overnight shaking at 37°C. From each overnight culture, 200 µL were plated on LA plates supplemented with 425 mg/L Pluronics and 100 mg/L rifampicin, and ten wells of a 96-well microtiter plate containing 100 µL LB supplemented with Pluronics and 16 mg/L rifampicin were inoculated with 2 µL culture each (in total 20 µL overnight culture) (Fig. 4G). All colonies on the plates containing 100 mg/L rifampicin were restreaked on the same selective plates and mutations within the rpoB gene were identified by local sequencing. From each positive well, 10 µL culture was plated on LA supplemented with Pluronics and 100 mg/L rifampicin and incubated overnight at 37°C. One colony per plate was restreaked on the same selective media followed by identification of mutations within the rpoB gene by local sequencing. For each of the five independent cultures, the types of mutations isolated from selections at 100 mg/L rifampicin were compared to the mutations isolated from selections using 16 mg/L rifampicin (Fig. 4I-M, table S3). The Null hypothesis for pre-existing mutations was that the mutations from 16 mg/L rifampicin should be a subset of the mutations from 100 mg/L rifampicin due to the ten-fold excess of culture used in the selection for high-level rifampicin resistance (Fig. 4H).

### PCR and local DNA sequencing

Polymerase chain reactions (PCR) for local DNA sequencing were performed using 2x PCR Master mix (Thermo Scientific). The following PCR program was used: 95°C for 3 min, 35 cycles of 95°C for 30 s, 50°C for 30 s, and 72°C for 1 min, followed by 72°C for 10 min. Local sequencing was carried out by Eurofins Genomics Europe Shared Services GmbH, Konstanz, Germany. Primer design and sequence analysis were performed using the CLC Main workbench 21.0.3 (CLCbio, QIAGEN, Denmark). See table S9 for a list of oligonucleotides.

### Microfluidics experiments

Microfluidic experiments were performed using a modified version of the PDMS mother machine microfluidic chip previously described (14). The version of the chip permitted simultaneous measurements of four different antibiotic concentrations (Fig. 1B). Medium pressure was controlled using a microfluidic flow controller (AstregoFlow).

For the microfluidic experiments, overnight cultures of cells were diluted 1,000-fold in fresh LB medium with Pluronics and grown shaking at 37°C for 4h before loading into the microfluidic chip. Cells were grown in the microfluidic chip for approximately 2h before the start of the experiments.

Microscopy was performed using a Ti2-E (Nikon) inverted microscope equipped with a CFI Plan Apochromat DM Lambda 1.45/100x objective (Nikon), a DMK 38UX304 camera (The Imaging Source), and a Spectra Gen. 1 (Lumencor) light source within a H201-ENCLOSURE hood with a H201-T-UNIT-BL controller (OKOlab). The microscope was controlled by Micro-Manager (version 1.4.23) (27) running in-house-built plugins. Phase-contrast images were acquired every minute or second minute with 80 ms exposure.

### Data analysis

Image analysis to determine single cell growth rates was done in MATLAB (Mathworks). Microscopy data were processed using an automated analysis pipeline developed previously (28), with one exception; segmentation of phase-contrast images was done using a U-net neural network (29) trained in-house. All raw data images and analysis software will be openly accessible through SciLifeLab data repository.

All other data analyses were performed with R 4.0.3 (30) and the results were plotted using the R package ggplot2 3.3.3 (31). Raw data and analyses scripts are available online.

## Acknowledgments

We thank S. Zikrin and O. Broström for help with the image analysis. This study was made possible by grants from the ERC (Advanced grant no. 885360), the Swedish Foundation for strategic Research (grant no. ARC19-0016), the Knut and Alice Wallenberg Foundation (grant no. 2016.0077, 2017.0291 and 2019.0439), and the eSSENCE e-science initiative. The computations and data management were enabled by resources provided by the Swedish National Infrastructure for Computing (SNIC) at UPPMAX, partially funded by the Swedish Research Council through grant agreement no. 2018-05973.

## Author contributions

Conceptualization: GB, JE

Experiments: GB, JL

Data analysis: GB

Funding acquisition: JE

Writing – original draft: GB

Writing – review & editing: GB, JL, JE

## Extended data

**Fig. S1.**
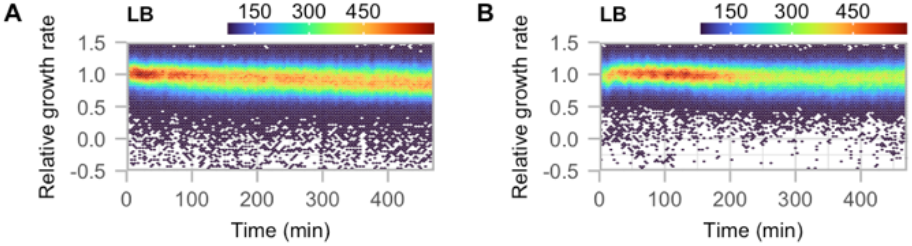
Single-cell growth rates during growth in LB. Distribution of single-cell growth rates as a function of time during the microfluidic experiments. The experiment was run in two biological replicates. The colors indicate the number of cells at each point.

**Fig. S2.**
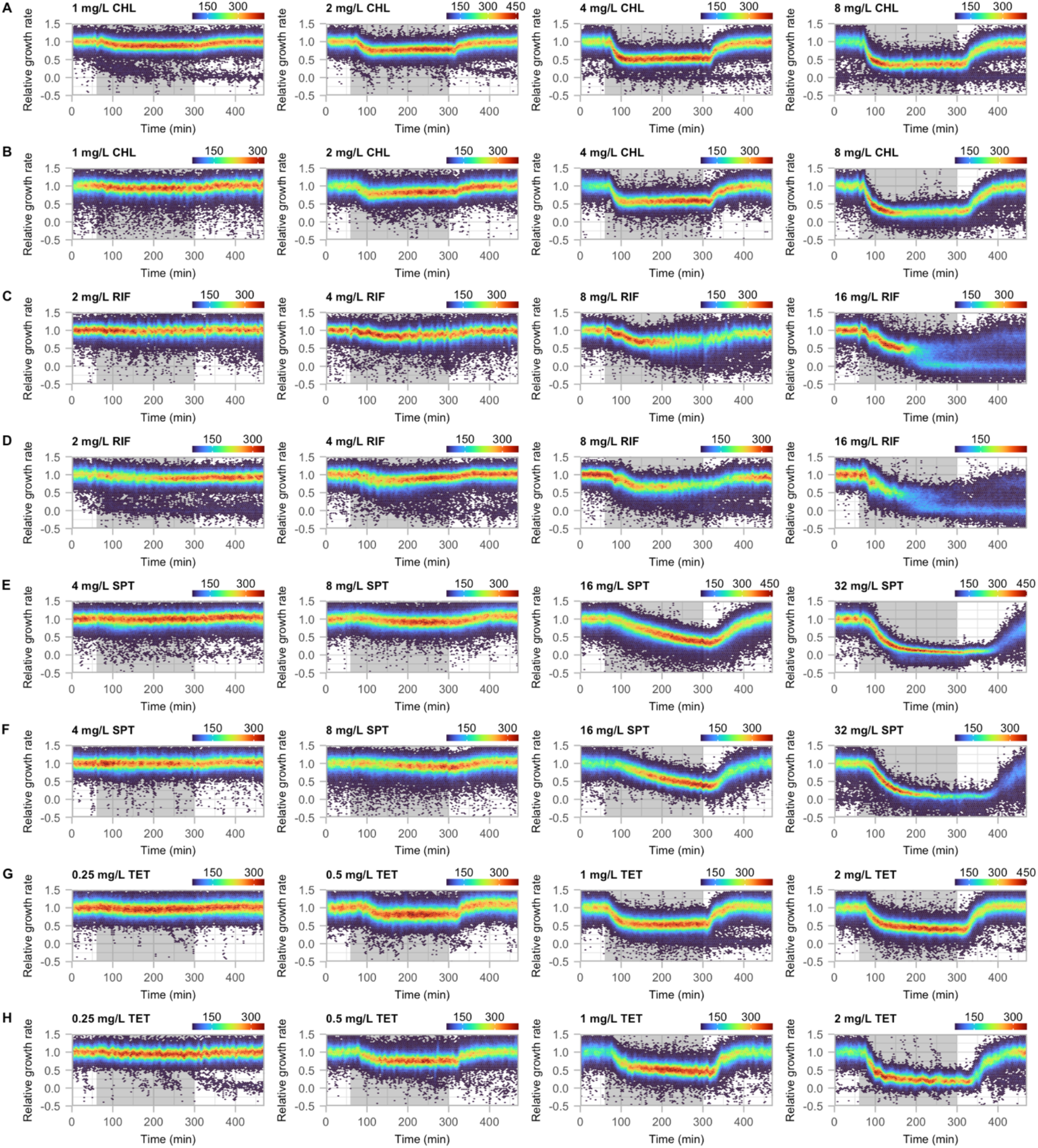
Single-cell growth rates during exposure to bacteriostatic antibiotics. Distribution of single-cell growth rates as a function of time during the microfluidic experiments. The antibiotic exposure phase is indicated by the grey background. The antibiotic concentrations correspond to 1/8x, 1/4x, 1/2x, and 1x MIC. Each antibiotic was run in two biological replicates. The colors indicate the number of cells at each point.

**Fig. S3.**
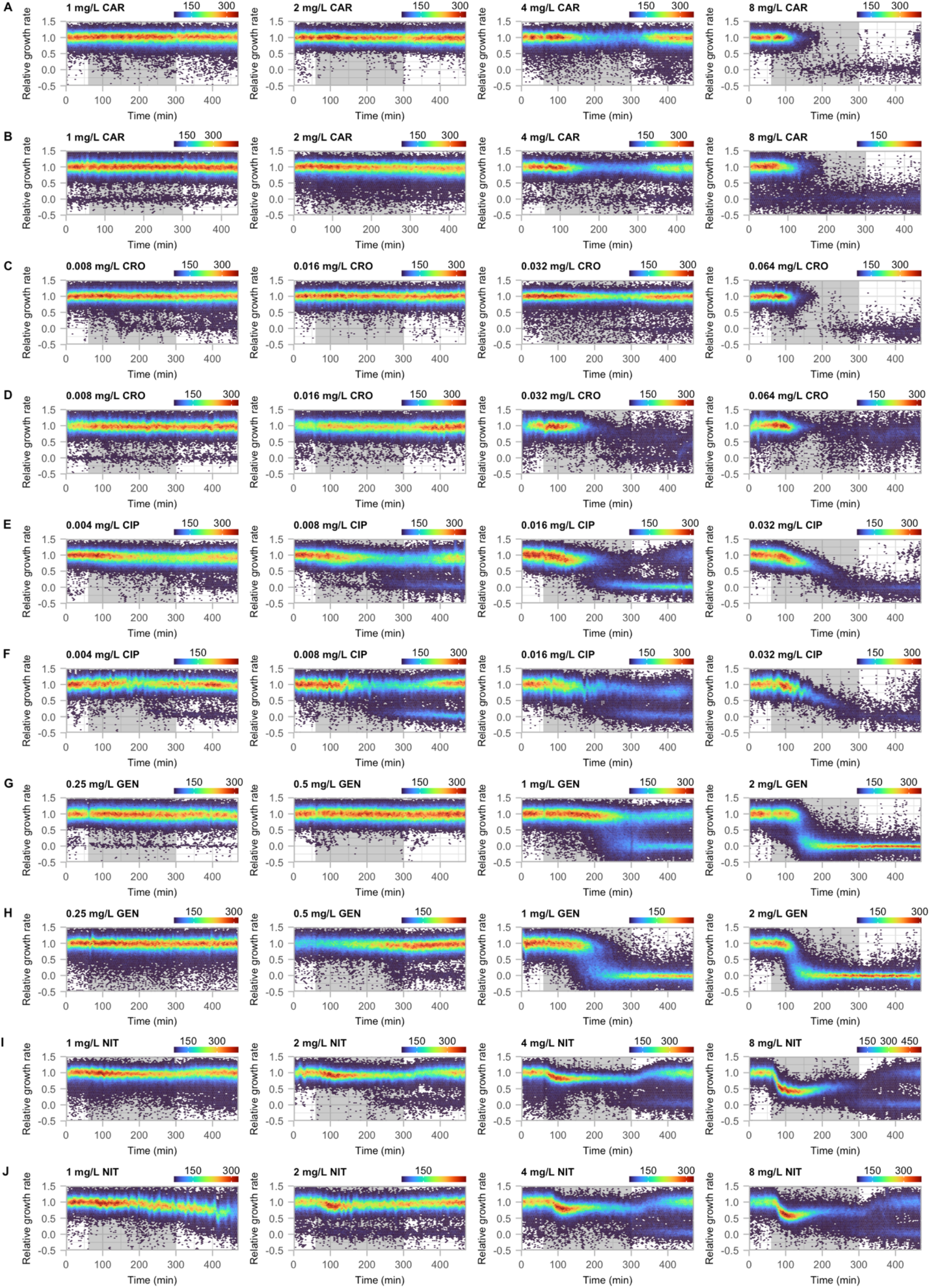
Single-cell growth rates during exposure to bactericidal antibiotics. Distribution of single-cell growth rates as a function of time during the microfluidic experiments. The antibiotic exposure phase is indicated by the grey background. The antibiotic concentrations correspond to 1/8x, 1/4x, 1/2x, and 1x MIC. Each antibiotic was run in two biological replicates. The colors indicate the number of cells at each point.

**Fig. S4.**
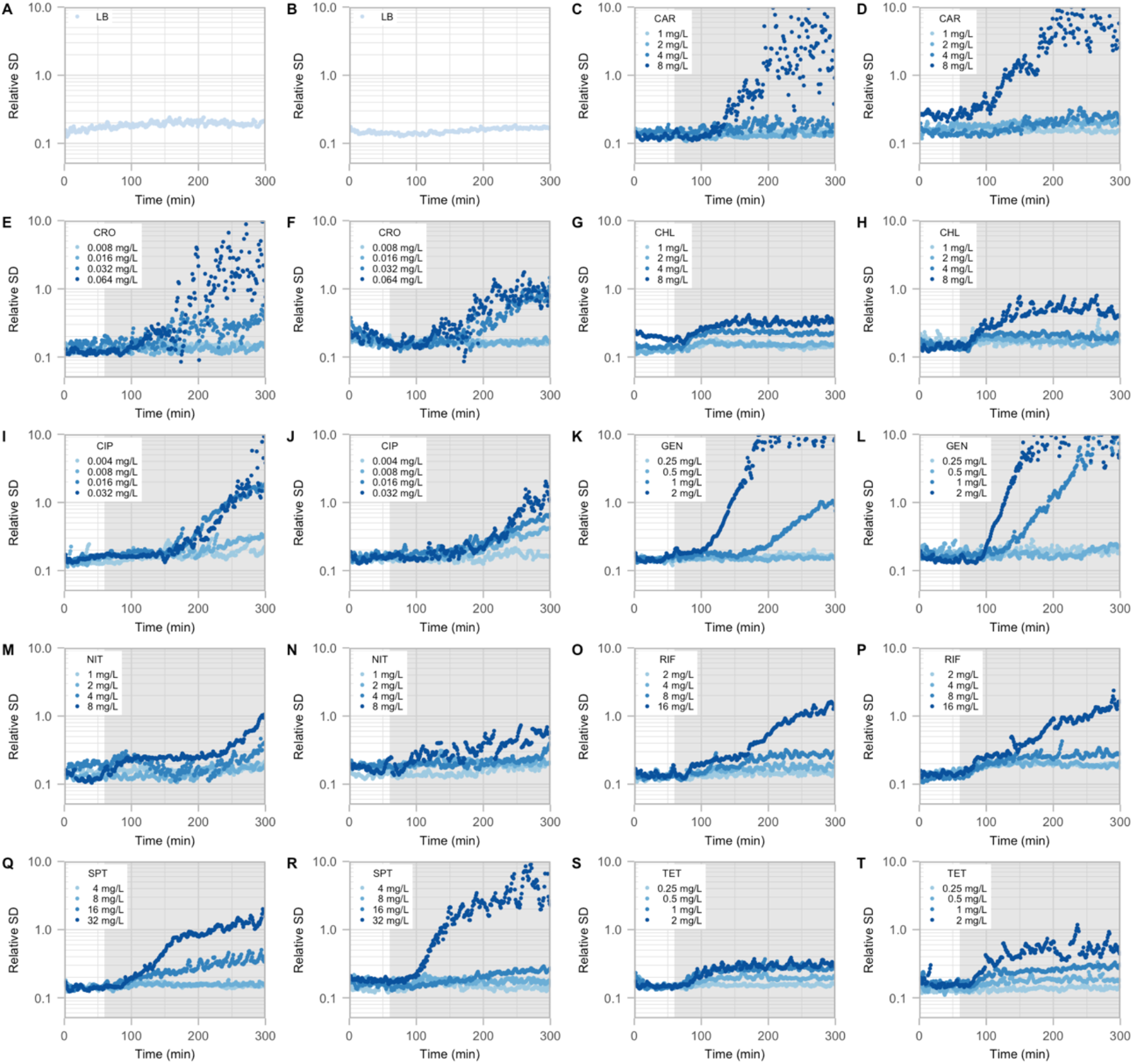
Cell-to-cell heterogeneity as function of time during antibiotic exposure. (A-T) Cell-to-cell heterogeneity, measured as relative standard deviation, as function of antibiotic exposure time. The period of antibiotic exposure is indicated by the grey background. Each antibiotic was run in two biological replicates.

**Fig. S5.**
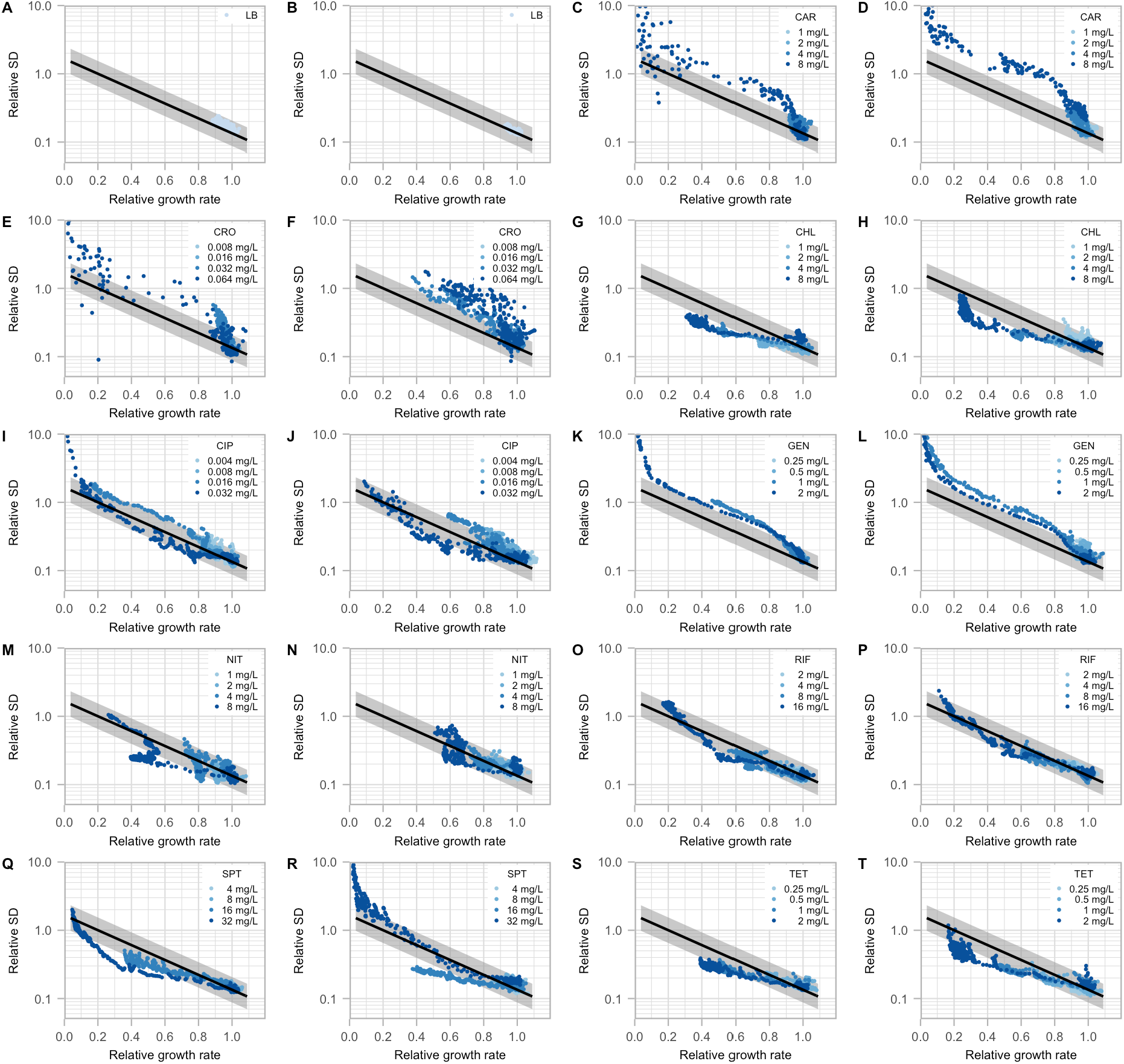
Cell-to-cell heterogeneity as function of growth rate during antibiotic exposure. (A-T) Cell-to-cell heterogeneity, measured as relative standard deviation, as function of relative growth rate. The black line and grey area indicate cell-to-cell heterogeneity that is expected from growth rate reduction (fig. S8). Each antibiotic was run in two biological replicates.

**Fig. S6.**
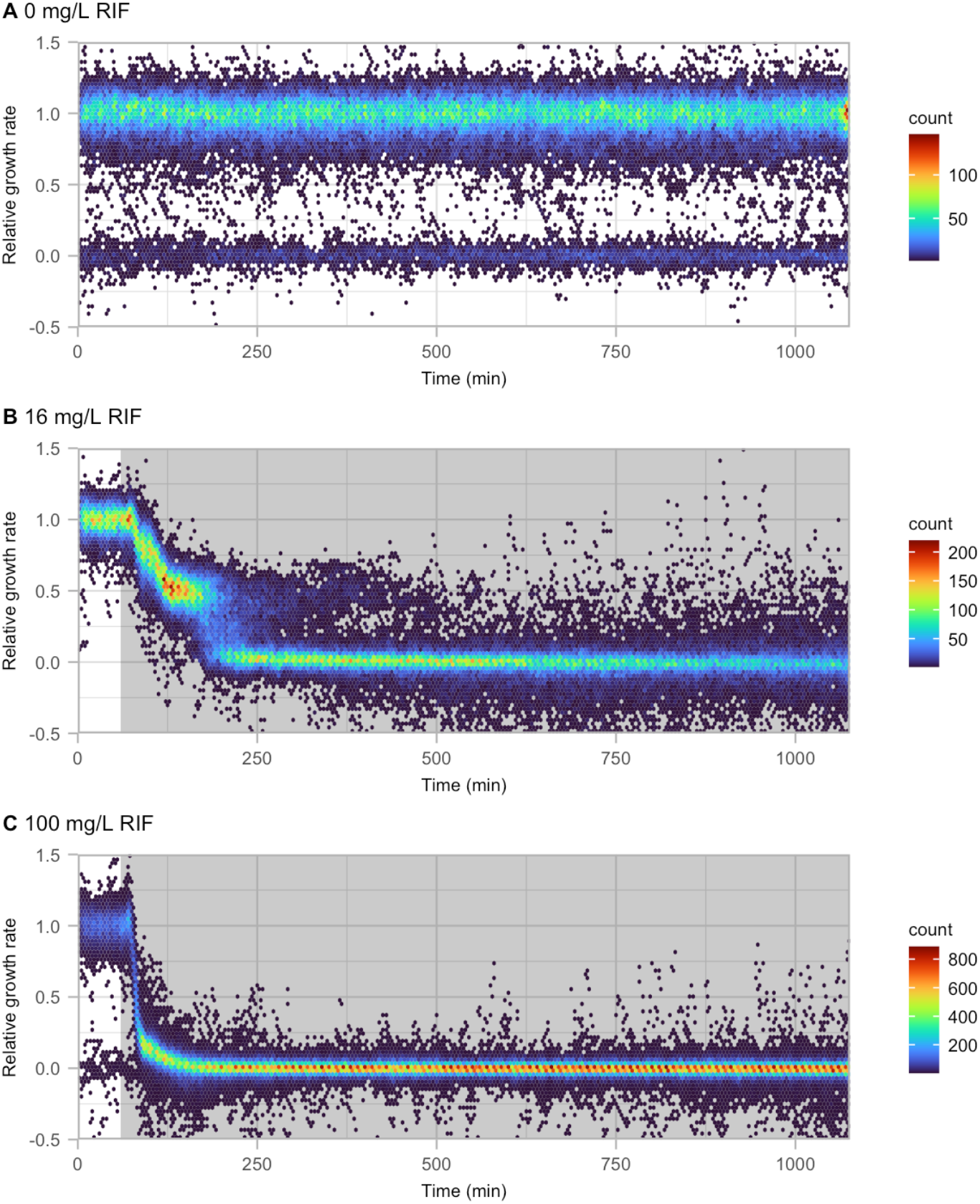
Single-cell growth rates of E. coli during 17h rifampicin exposure. Distribution of single-cell growth rates of E. coli as a function of time during the microfluidic experiment with 17h exposure to (A) LB, (B), 1x MIC rifampicin, and (C) ∼6x MIC rifampicin. The antibiotic exposure phase is indicated by the grey background. The colors indicate the number of cells at each point.

**Fig. S7.**
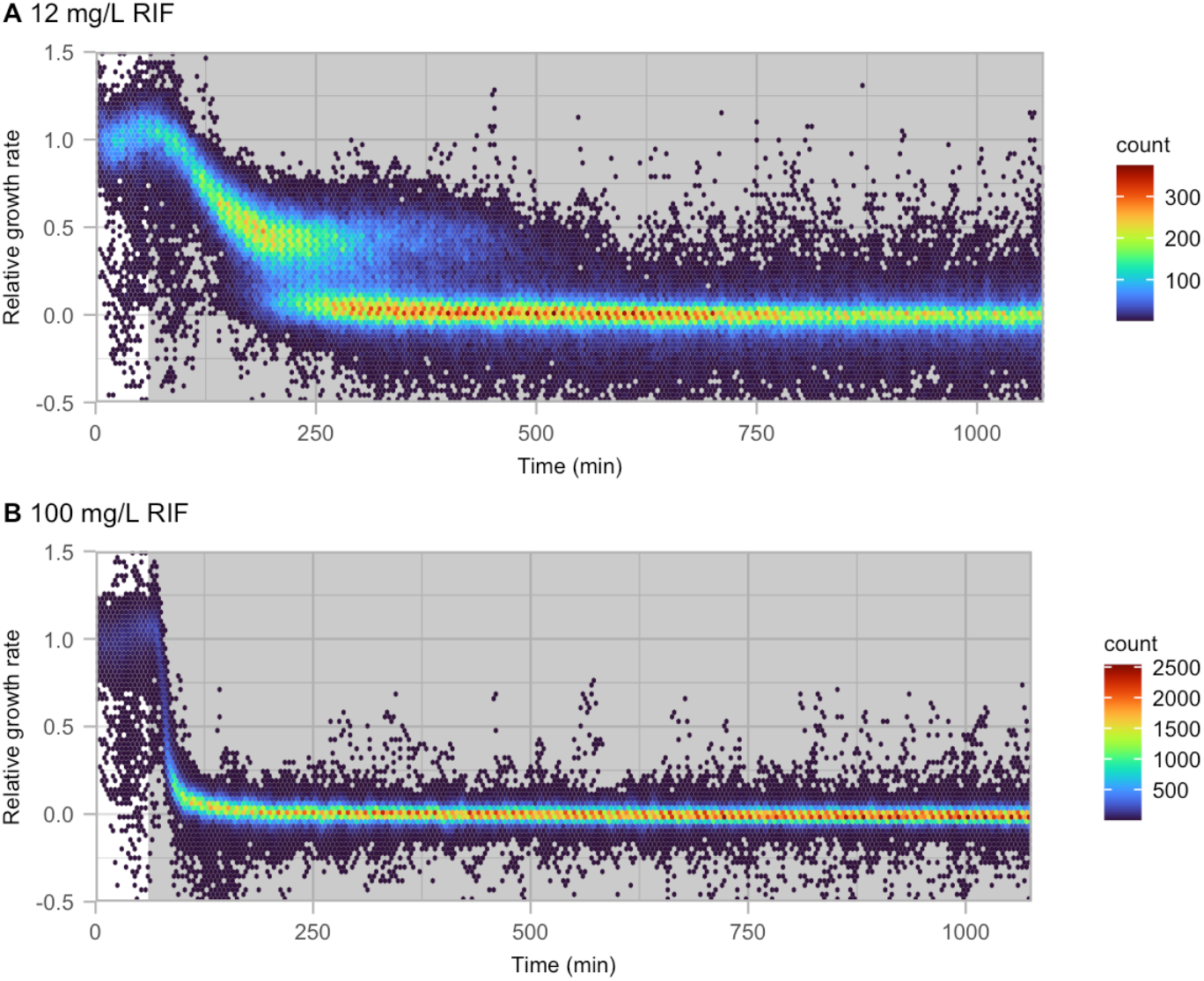
Single-cell growth rates of S. enterica during 17h rifampicin exposure. Distribution of single-cell growth rates of S. enterica as a function of time during the microfluidic experiment with 17h exposure to (A) LB, (B), 1x MIC rifampicin, and (C) ∼6x MIC rifampicin. The antibiotic exposure phase is indicated by the grey background. The colors indicate the number of cells at each point.

**Fig. S8.**
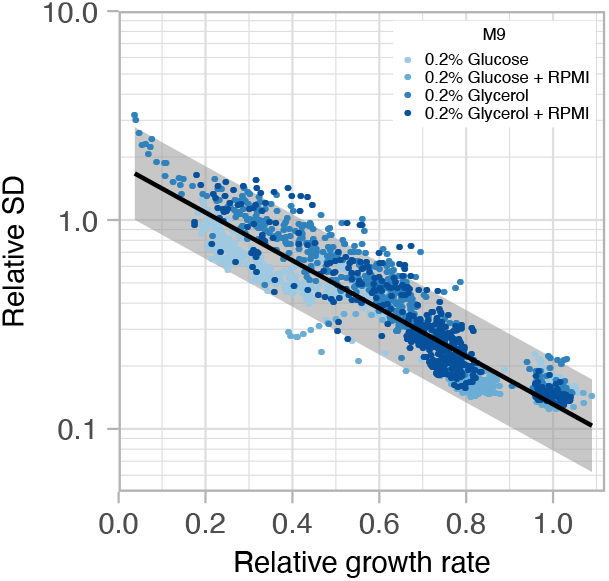
Cell-to-cell heterogeneity as function of growth rate in minimal media. Cell-to-cell heterogeneity, measured as relative standard deviation, as function of relative growth rate after switching to growth in minimal media. The black line corresponds to the modeled trend line and the grey area includes 95% of all data points.

**Fig. S9.**
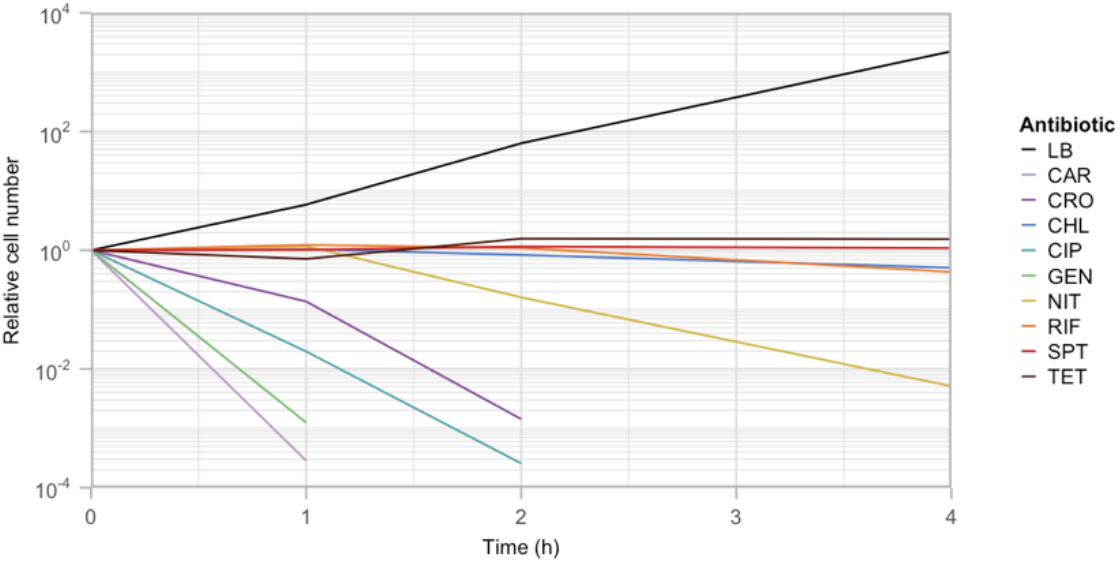
Time-kill kinetic assay for the antibiotics used in the study. Relative cell number as function of antibiotic exposure time. Each antibiotic was added at a concentration corresponding to 8x MIC.

**Table S1.** Overview of antibiotics used in the study.

| **Antimicrobial** | **Drug class** | **Drug target(s)** | **Type of activity<sup>a</sup>** |
| --- | --- | --- | --- |
| Carbenicillin (CAR) | β-lactam | Cell wall synthesis | Bactericidal |
| Ceftriaxone (CRO) | Cephalosporin | Cell wall synthesis | Bactericidal |
| Chloramphenicol (CHL) | Amphenicol | Protein synthesis (50S) | Bacteriostatic |
| Ciprofloxacin (CIP) | Fluoroquinolone | DNA replication / cell division | Bactericidal |
| Gentamycin (GEN) | Aminoglycoside | Protein synthesis (30S) | Bactericidal |
| Nitrofurantoin (NIT) | Nitrofuran | Multiple | Bactericidal |
| Rifampicin (RIF) | Rifamycin | Transcription | Bacteriostatic |
| Spectinomycin (SPT) | Aminoglycoside | Protein synthesis (30S) | Bacteriostatic |
| Tetracycline (TET) | Tetracycline | Protein synthesis (30S) | Bacteriostatic |

<sup>a</sup> Bactericidal and bacteriostatic effects for the conditions used in this study were tested using a time-kill kinetic assays (Fig. S9).

**Table S2.** Overview of antibiotic concentrations used for *E. coli* in the study. Antibiotic Concentration (mg/L)

| **Antibiotic** | **Concentration (mg/L)** |  |  |  |
| --- | --- | --- | --- | --- |
|  | **1/8x MIC** | **1/4x MIC** | **1/2x MIC** | **1x MIC** |
| Carbenicillin | 1 | 2 | 4 | 8 |
| Ceftriaxone | 0.008 | 0.016 | 0.032 | 0.064 |
| Chloramphenicol | 1 | 2 | 4 | 8 |
| Ciprofloxacin | 0.004 | 0.008 | 0.016 | 0.032 |
| Gentamycin | 0.25 | 0.5 | 1 | 2 |
| Nitrofurantoin | 1 | 2 | 4 | 8 |
| Rifampicin | 2 | 4 | 8 | 16 |
| Spectinomycin | 4 | 8 | 16 | 32 |
| Tetracycline | 0.25 | 0.5 | 1 | 2 |

**Table S3.** Overview of *rpoB* mutations selected in this study.

| Culture | Panel<sup>a</sup> | Unique *rpoB* mutations identified during selection with |  |
| --- | --- | --- | --- |
|  |  | 16 mg/L RIF | 100 mg/L RIF |
| 1 | I | Q513L, S531insVS, L533H,<br>L533P, P564L | V146F, D516G, H526L, L533H,<br>I572F |
| 2 | J | S512P, Q513P, D516G, H526Y,<br>I572F | D6516G |
| 3 | K | D516N, L533P | H526Y |
| 4 | L | Q513R, D516G, I572F | Q513P, D516Y |
| 5 | M | S512Y | S531F |

<sup>a</sup> Underlying data for the Venn-diagrams in the corresponding panels of fig 4.

**Table S4.** Antibiotic concentrations used to select for resistance in various species.

| Species | Antibiotic | Antibiotic concentration during resistance selection |  |
| --- | --- | --- | --- |
|  |  | low | high |
| *E. coli* | RIF | 16 mg/L | 100 mg/L |
| *E. coli* | NIT | 8 mg/L | 32 mg/L |
| *S. enterica* | RIF | 8 mg/L | 100 mg/L |
| *K. pneumoniae* | RIF | 16 mg/L | 100 mg/L |
| *S. aureus* | RIF | 0.0032 mg/L | 0.1 mg/L |
| *E. faecalis* | RIF | 1 mg/L | 10 mg/L |

**Table S5.**
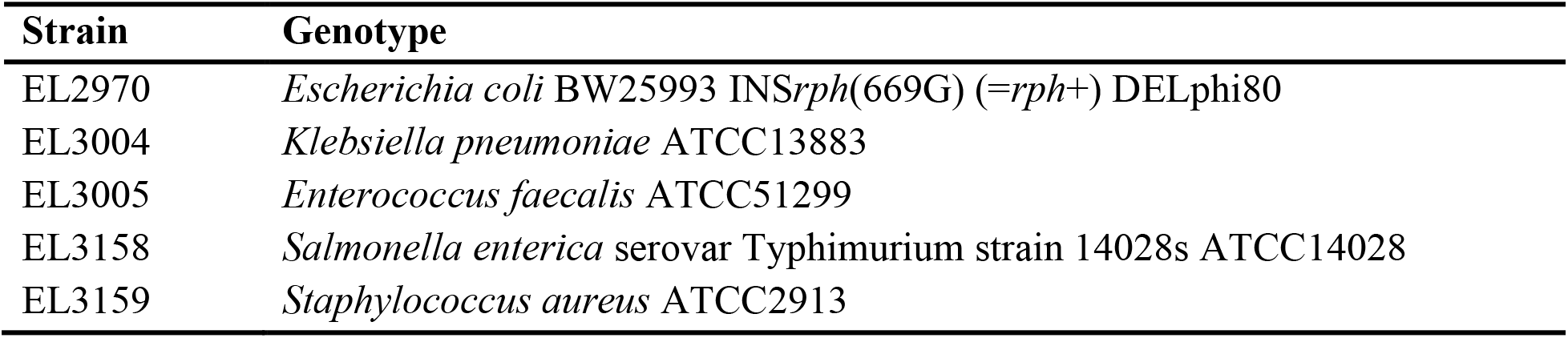
Strain list.

| **Strain** | **Genotype** |
| --- | --- |
| EL2970 | *Escherichia coli* BW25993 INSrph(669G) (=rph+) DELphi80 |
| EL3004 | *Klebsiella pneumoniae* ATCC13883 |
| EL3005 | *Enterococcus faecalis* ATCC51299 |
| EL3158 | *Salmonella enterica* serovar Typhimurium strain 14028s ATCC14028 |
| EL3159 | *Staphylococcus aureus* ATCC2913 |

**Table S6.** Tested concentration range and results of MIC tests.

| Antibiotic | Species | Tested concentration range<sup>a</sup> | MIC<sup>b</sup> |
| --- | --- | --- | --- |
| CAR | *Eco* | 0.5 – 32 mg/L | 8 mg/L |
| CRO | *Eco* | 0.002 – 0.128 mg/L | 0.064 mg/L |
| CHL | *Eco* | 0.25 – 16 mg/L | 8 mg/L |
| CIP | *Eco* | 0.001 – 0.064 | 0.032 mg/L |
| GEN | *Eco* | 0.125 – 8 mg/L | 2 mg/L |
| RIF | *Eco* | 1 – 16 mg/L | 16 mg/L |
| RIF | *Sty* | 0.5 – 32 mg/L | 8 mg/L |
| RIF | *Kpn* | 4 – 256 mg/L | 16 mg/L |
| RIF | *Sau* | 0.001 – 0.0064 mg/L | 0.0032 mg/L |
| RIF | *Efa* | 0.5 – 32 mg/L | 1 mg/L |
| NIT | *Eco* | 0.5 – 32 mg/L | 8 mg/L |
| SPT | *Eco* | 1 – 64 mg/L | 32 mg/L |
| TET | *Eco* | 0.125 – 8 mg/L | 2 mg/L |

<sup>a</sup> The MIC tests were performed using a two-fold dilution series between the two indicated values.
<sup>b</sup> Result of three biological replicates.

**Table S7.**
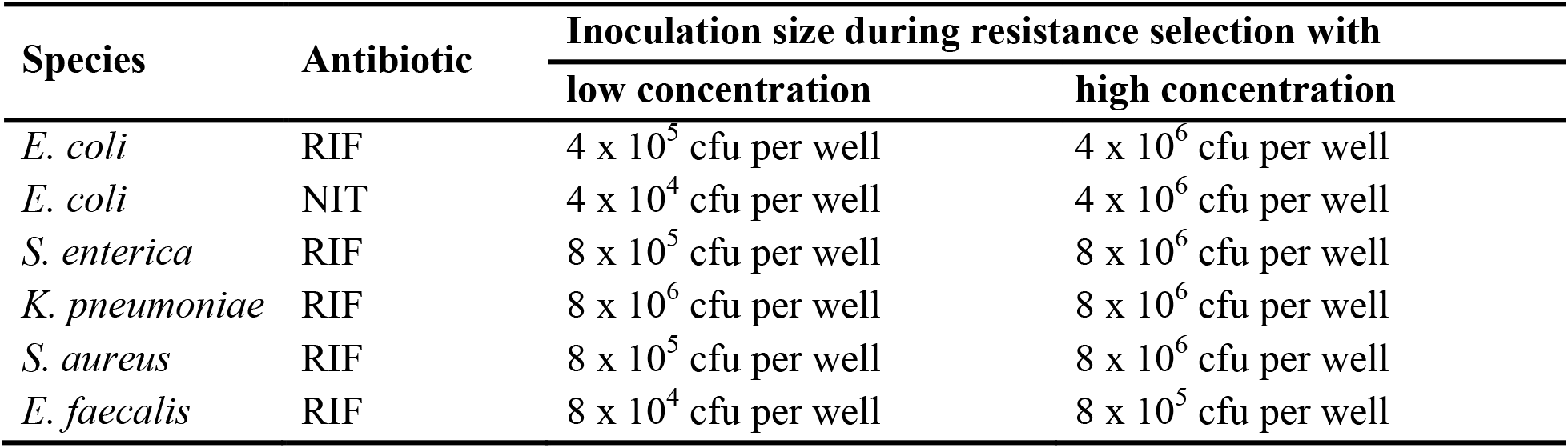
Inoculation sizes used to select for antibiotic resistance in various species.

**Table S8.** Results from intrinsic mutation rate determination.

| **RpoB allele** | **Selected in<sup>a</sup>** | **Positive/negative cultures** | **$\mu^{100\text{STR}}$ <sup>b</sup>** | **P value<sup>c</sup>** |
| --- | --- | --- | --- | --- |
| Wild-type | - | 2/18 | $3.76 \times 10^{-11}$ | - |
| D516G | 16 mg/L | 3/17 | $5.80 \times 10^{-11}$ | 0.46 |
| D516G | 100 mg/L | 2/18 | $3.76 \times 10^{-11}$ | 1 |
| H526Y | 16 mg/L | 2/18 | $3.76 \times 10^{-11}$ | 1 |
| H526Y | 100 mg/L | 1/19 | $1.83 \times 10^{-11}$ | 0.46 |
| I572F | 16 mg/L | 3/17 | $5.80 \times 10^{-11}$ | 0.46 |
| I572F | 100 mg/L | 4/16 | $7.97 \times 10^{-11}$ | 0.14 |

<sup>a</sup> Rifampicin concentration that the *rpoB* mutation initially was selected in.
<sup>b</sup> Mutation rate for resistance to 100 mg/L streptomycin (per cell per generation).
<sup>c</sup> Chi-squared test comparing the number of positive and negative cultures of the mutants with those of the wild-type strain.

**Table S9.**
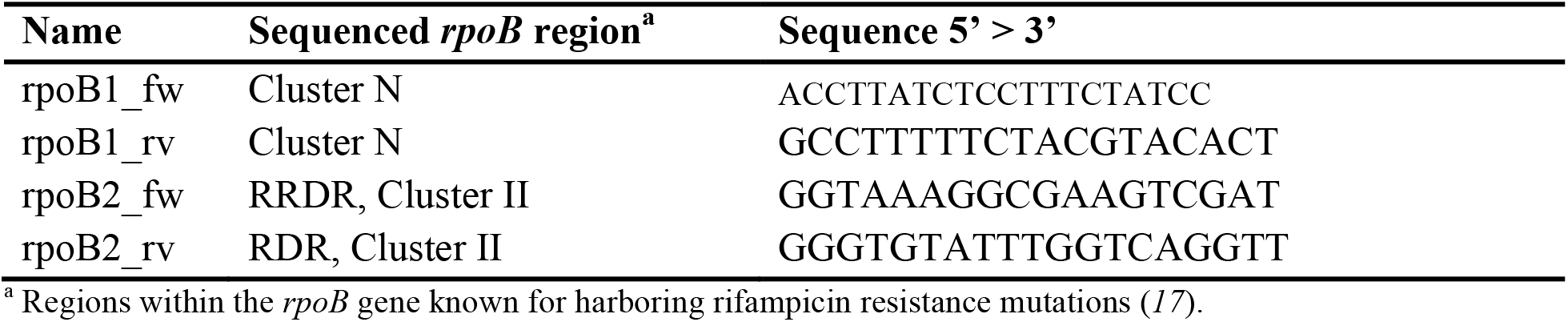
List of oligonucleotides used to identify *rpoB* mutations.

## Supplementary information

<u>Movie S1:</u> (Top) Representative video of the microfluidic growth experiments. E. coli cells were grown for 1h in LB with Pluronics. Then the media supply was changed to fresh LB with Pluronics (A) without rifampicin, or (B-E) with rifampicin corresponding to 1/8x, 1/4x, 1/2x, or 1x MIC. After 4 h, the media supply was changed back to LB with Pluronics for 3 h. The media change is indicated by the background color (green: LB, red: RIF). Phase contrast images were taken at intervals of 1 min. The playback rate of the video is 10 fps. (Bottom) Distribution of single-cell growth rates as a function of time during the microfluidic experiments.

